# Direct nanopore sequencing of *M. tuberculosis* on sputa and rescue of suboptimal results to enhance transmission surveillance

**DOI:** 10.1101/2025.09.23.678181

**Authors:** Sheri M. Saleeb, Andrea Marcos-Abellán, María Teresa Cabezas Fernández, Silvia Vallejo-Godoy, Miguel Martínez-Lirola, Guadalupe Bernal Ramirez, Marta Herranz-Martín, Sergio Buenestado-Serrano, Alfonso Pardo-Diaz, María Jesús Ruiz Serrano, Patricia Muñoz, Laura Pérez-Lago, Darío García de Viedma

**Affiliations:** Department of Clinical Microbiology and Infectious Diseases, Gregorio Marañón General University Hospital. Gregorio Marañón Health Research Institute (IiSGM), Madrid, Spain; CIBER of Enfermedades respiratorias (CIBERES), Spain; Doctoral school, Autonomous University of Madrid, Spain; Microbiology department. Torrecárdenas University Hospital Complex, Almería, Spain; Department of Preventive Medicine, Public Health and Epidemiological Surveillance, Poniente University Hospital, Almería, Spain; Department of Technology, Extremadura Center for Advanced Technologies (CETA-CIEMAT), Trujillo, Spain; Department of Medicine, Complutense University of Madrid, Spain

**Keywords:** Tuberculosis, transmission, genomic epidemiology, nanopore sequencing, direct on sputa

## Abstract

Whole-genome sequencing (WGS) enhances precision in predicting antimicrobial resistance and tracking *Mycobacterium tuberculosis* (MTB) transmission. Due to MTB’s slow-growing nature, genomic results are delayed; however, few efforts have sought to accelerate them by performing WGS directly on respiratory specimens. Most culture-free efforts have focused on accelerating resistance prediction. The present study provides further evidence to the only preceding study aiming to accelerate precise delineation of transmission, coupling culture-free WGS to a surveillance programme. Our study is distinguished from its predecessor by being the first to apply flexible nanopore sequencing to further accelerate the process. A total of 71 sputa were selected, in which we applied only a procedure to deplete human DNA, thus avoiding costly and cumbersome capture-bait alternatives. Optimal results (>90% genome covered, mean coverage >45× and >70% genome covered >20×) were obtained from 33.8% of cases, allowing the assignment to transmission clusters close to diagnosis of every new case. A further 12.6% of samples yielded suboptimal results (15.5%–90.92% at >10×), which were exploited through a rescue pipeline. This approach was based on identifying informative SNPs acting as markers for relevant transmission clusters in our population. The pipeline enabled pre-allocation of new cases to pre-existing clusters and, in some cases, precise genomic relationships with the preceding cases in the cluster. In summary, this study demonstrates that epidemiologically valuable information can be obtained directly from sputum in approximately half the samples analysed. It represents a new advancement in the pursuit of faster comparative genomics, with epidemiological purposes, at diagnosis.

## Introduction

Whole-genome sequencing (WGS) has significantly enhanced the precision with which we can track *Mycobacterium tuberculosis* (MTB) transmission, thereby optimising the responses required for transmission control and guiding epidemiological interventions (1). However, the implementation of these interventions necessitates the minimisation of the time interval between the diagnosis of each new case and the availability of genomic epidemiology analyses. The majority of genomic analysis conducted on MTB for epidemiological purposes is performed on cultures, a process that introduces significant delays due to the slow growth rate of MTB (2). Consequently, the optimal approach to expedite response times in genomic epidemiology entails a direct transition from cultures to sequencing on clinical specimens, thereby circumventing the culture step entirely to facilitate enhanced epidemiological decision-making. Despite the endeavours to perform WGS directly on sputa (3–6), these are still limited, with several requiring either demanding long procedures (4) or enrichment by capturing MTB material (2,3,5). All of the aforementioned aspects indicate that WGS analysis on sputa remains challenging (7). Only targeted approaches, devoted to MTB identification, resistance mutations, or general phylogenetic analysis, have been demonstrated to be successful (8–11).

In the present study, the performance of fast nanopore sequencing was evaluated directly on sputa, without the need for additional capture/enrichment procedures, within the context of our long-term genomic epidemiology surveillance/intervention program in Almeria (12). This approach was adopted to ensure a faster analysis, with the objective of tailoring epidemiological interventions. Furthermore, an alternative rescue pipeline was developed to exploit suboptimal sequences and assign them to preexisting relevant clusters by focusing on cluster marker SNPs. The present study constitutes a further progression in the direction of real-time genomic surveillance in routine laboratory settings.

## Materials and Methods

### Sample collection

Clinical samples (decontaminated sputa with a bacillary load ranging, according to smear grade, from +1 to +4) were routinely collected at Torrecárdenas University Hospital (Almería, Spain). The samples were then decontaminated using a solution of N-acetyl-L-cysteine and NaOH (final NaOH concentration: 1%), followed by pH neutralisation with 10–15 mL of phosphate buffer (pH 6.8) for resuspension. The samples were then transferred frozen at −20 °C to our laboratory at Gregorio Marañón Hospital (Madrid, Spain).

### DNA purification

Ten to fifteen millilitres of decontaminated sputa were subjected to a centrifugal process at 8,500 revolutions per minute for 40 minutes at a temperature of 4°C. Thereafter, the supernatant was discarded, and the resultant pellet was suspended in 500 microlitres of AHL buffer. The depletion of human DNA, extraction, and purification, were performed by the QIAamp DNA Microbiome Kit (Qiagen, Hilden, Germany), following the manufacturer’s instructions. The concentration and purification of DNA was achieved by employing 1.8X magnetic beads (CleanNGS, GC biotech, Netherlands) and eluting in 12 µl nuclease-free water. The quantification of the purified DNA was conducted using a Qubit fluorometer.

### Real-time PCR

In order to ascertain the relative proportion of human, bacterial, and *Mycobacterium tuberculosis* complex (MTBC) DNA in the purified DNA samples, three real-time PCRs were performed using TaqMan probes to target, respectively, *RNase P*, *16S rRNA*, and *Rv2341* (13) genes, using the following primers/probes: *RNase P* Forward-AGATTTGGACCTGCGAGCG, Reverse-GAGCGGCTGTCTCCACAAGT, Probe-HEX-TTCTGACCTGAAGGCTCTGCGCGBHQ-1; *16S rRNA*: Forward 5’-TGGAGCATGTGGTTTAATTCGA-3’, Reverse 5’-TGCGGGACTTAACCCAACA-3’, probe 5’-HEX-CACGAGCTGACGACARCCATGCA-3’; for *Rv2341* Forward 5’-GGCCGCTCATGCTCCTTGGAT-3’, reverse 5’-AGGTCGGTTCGCTGGTCTTG-3’, and probe 5’-6-FAM-TGAGTGCCTGCGGCCGCAGCGC-BHQ-1-3’. The PCR reactions incorporated 0.25 µM of each primer pair and probe, with 2 µL of the LightCycler FastStart DNA Master HybProbe (Roche, USA) buffer, containing 3.2 mM Mg2+, in a final reaction volume of 20 µL. The PCR conditions were for *RNase P* 55°C for 5 minutes, 95°C for 5 minutes, followed by 35 cycles of 95°C for 15 seconds, and at 60°C for 30 seconds, concluding with 40°C for 30 seconds; for *16S rRNA*, 95°C for 10 minutes, 40 cycles of 95°C for 10 seconds, 60°C for 20 seconds, 72°C for 1 second, and finally 40°C for 30 seconds; for *Rv4321*, 95°C for 10 minutes, 55 cycles for 95°C for 10 seconds, 60°C for 20 seconds, 62°C for 1 minute and finally, 40°C in 30 seconds.

### Whole genome sequencing

Libraries were prepared from all the purified DNA obtained from each sputum, using the Rapid PCR Barcoding kit (SQK-RPB004 & SQK-RPB114) as per the manufacturer’s instructions (Oxford Nanopore Technologies, Oxford, UK). 5% Dimethyl sulfoxide (DMSO) was added to improve DNA denaturation during the amplification step. As recommended in the first step of the library preparation protocol (fragmentation step), 5 ng of DNA should be adjusted in 3 µl. However, since the amount of DNA obtained from sputa was below these values, 2-3 libraries were prepared per specimen, to be later pooled. The libraries were loaded and run on MinION flow-cells (R.9.4.1 FLO-MIN106 & R10.4.1 FLOW-MIN114). Genome coverage and depth were monitored during the sequencing run using RAMPART v1.2.0. The sequencing run was permitted to continue until achieving a minimum of 70% genome coverage at 20× depth in all the sequences, or until 72 hours had elapsed, whichever occurred first. Furthermore, when the progression of the run indicated that the required coverage thresholds were not going to be obtained for some of the samples, the run was aborted to allow the flow-cell to retain sufficient pores for reuse in other projects.

### Bioinformatic analysis

#### Standard pipeline

The subsequent analysis of the data was conducted using an in-house pipeline (GitHub: MG-IiSGM/prokaION) (14). Initially, the raw fast5/pod5 files underwent a preprocessing step, whereby they were converted into fastq format through base-calling and barcoding using Guppy v6.4.6, or Dorado v0.9.6. Subsequently, the quality of the data was assessed using NanoFilt v2.8.0 and NanoPlot v1.41.0, in order to ensure the reliability and informativeness of the data for subsequent analyses. The resulting reads were then mapped using a hypothetical MTB ancestral genome (15) as a reference, with minimap2 v2.28 employed for this purpose. Subsequently, single-nucleotide polymorphisms (SNPs) were identified with FreeBayes v1.3.2. The detected variants were subsequently annotated with SnpEff v5.1, and occasional low-coverage positions (>12×) were recalibrated using joint variant calling. The identification of human and bacterial samples was conducted utilising the Kraken2 2.1.3 and Mash v2.3 software.

For the assignment of new cases to clusters we introduced the new sequences into the global phylogeny obtained for the 1,173 cases sequenced in Almería. In instances where a new sequence was found to be in close proximity to a preexisting sequence, either within the same clade or as a sister taxon originating from the same rooted branch, all related sequences were processed to generate a pairwise SNP distance matrix. A threshold of 12 SNP differences was applied: sequences within this limit were considered part of the same cluster, whereas sequences exceeding this threshold were classified as outgroup “orphans”.

The alignment and SNPs were then visualised and inspected using the IGV (Integrative Genomics Viewer) program. Median-joining networks of genome-related isolates were constructed from the SNP matrix generated by PopART software (16).

#### Rescue pipeline to exploit suboptimal sequences

Firstly, the identification of set of marker SNPs for a selection of relevant clusters (those growing in the 2021-25 period) was undertaken, to ascertain the capacity of these informative marker SNPs to be called from suboptimal sequences. The process involved, firstly, the identification of common SNPs shared by all the representatives of the relevant clusters that had been selected. Then, cluster marker SNPs were identified by filtering cluster common SNPs against an in-house global reference database comprising SNPs from 109,648 high-quality sequences from publicly available datasets, representing the geographic and phylogenetic diversity of MTB lineages 1–9. The inclusion of sequences was contingent upon the fulfilment of stringent selection criteria, encompassing the following criteria: only datasets obtained by Illumina, the presence of complete metadata, exclusive taxonomic assignment to MTB, and the attainment of specified quality thresholds (>70% of the genome covered at >20× depth and <25% of the genome uncovered). The global SNPs database contains a total of 2,042,025 polymorphic positions (allele depth >20, allele frequency >70%). The final set of marker SNPs for the clusters of interest was subsequently utilised in a rescue pipeline designed to exploit suboptimal sequences. In instances where more than one marker SNP is found within the same gene, they are not taken into consideration, as it is deemed to be an extremely improbable occurrence in MTB and it may suggest false calls.

Suboptimal sequences were preassigned as likely to correspond to one of the relevant clusters, if by applying the rescue pipeline, >25% of the marker SNPs of a one of the relevant clusters were called at ≥ 5x.

#### Epidemiological data collection

The Almería TB Prevention and Control Program (TBPCP) obtained and analysed epidemiological data. The TBPCP involves epidemiologists, microbiologists, professionals in preventive medicine units, nurses, social workers, professionals in primary care centres and three hospital tuberculosis clinical units. The TBPCP is integrated into the Epidemiological Surveillance System of Andalusia (SVEA), an extensive surveillance system integrated within the Spanish Epidemiological Surveillance Network (RENAVE). Community health workers (CHWs) belonging to local non-governmental organisations (Red Cross and the Consortium of Entities for Comprehensive Action with Migrants) operating within the designated territory were involved, to provide support to the public health system and to serve as cultural mediators and translators.

A data collection form was used to record the main cluster characteristics, transmission environments and risk factors identified. To supplement the findings, a comprehensive review of the patients’ clinical records and additional interviews with were conducted for the cases in clusters that required a more profound investigation.

## Results

### General performance of nanopore sequencing from sputa

The analysis encompassed 71 stain-positive sputa from different patients. Forty-one of these corresponded to prospective non-selected consecutive cases, diagnosed in the period January – October 2024. The remaining cases were analysed retrospectively, in two different periods, including sputa corresponding to the years 2007-2015 and November 2024-June 2025, respectively (Table 1). A total of 25 nanopore sequencing runs were conducted, incorporating libraries for 1–5 specimens per run (median 3 samples; 4.9–6.9 fmol/sample).

**Table 1:**
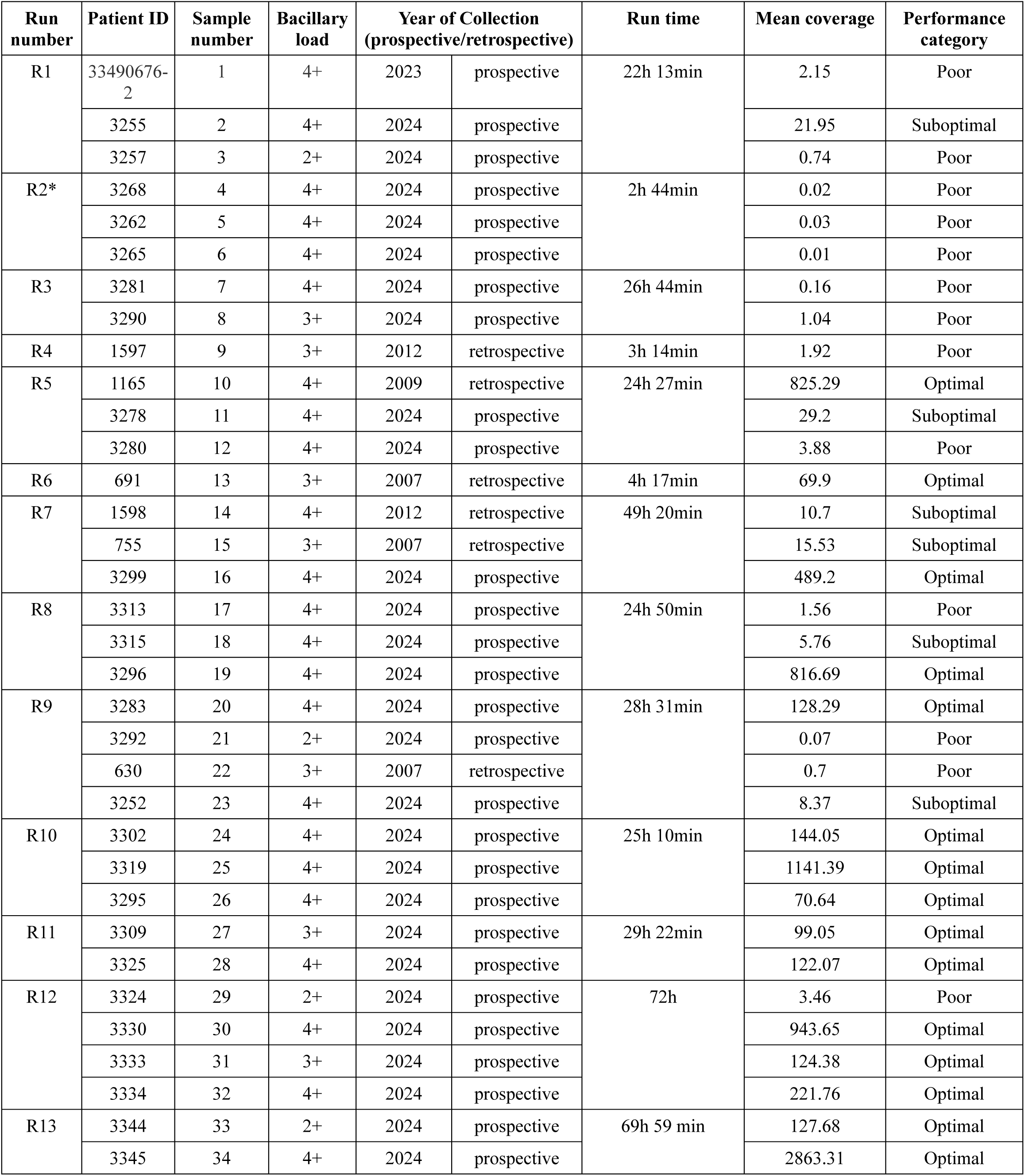
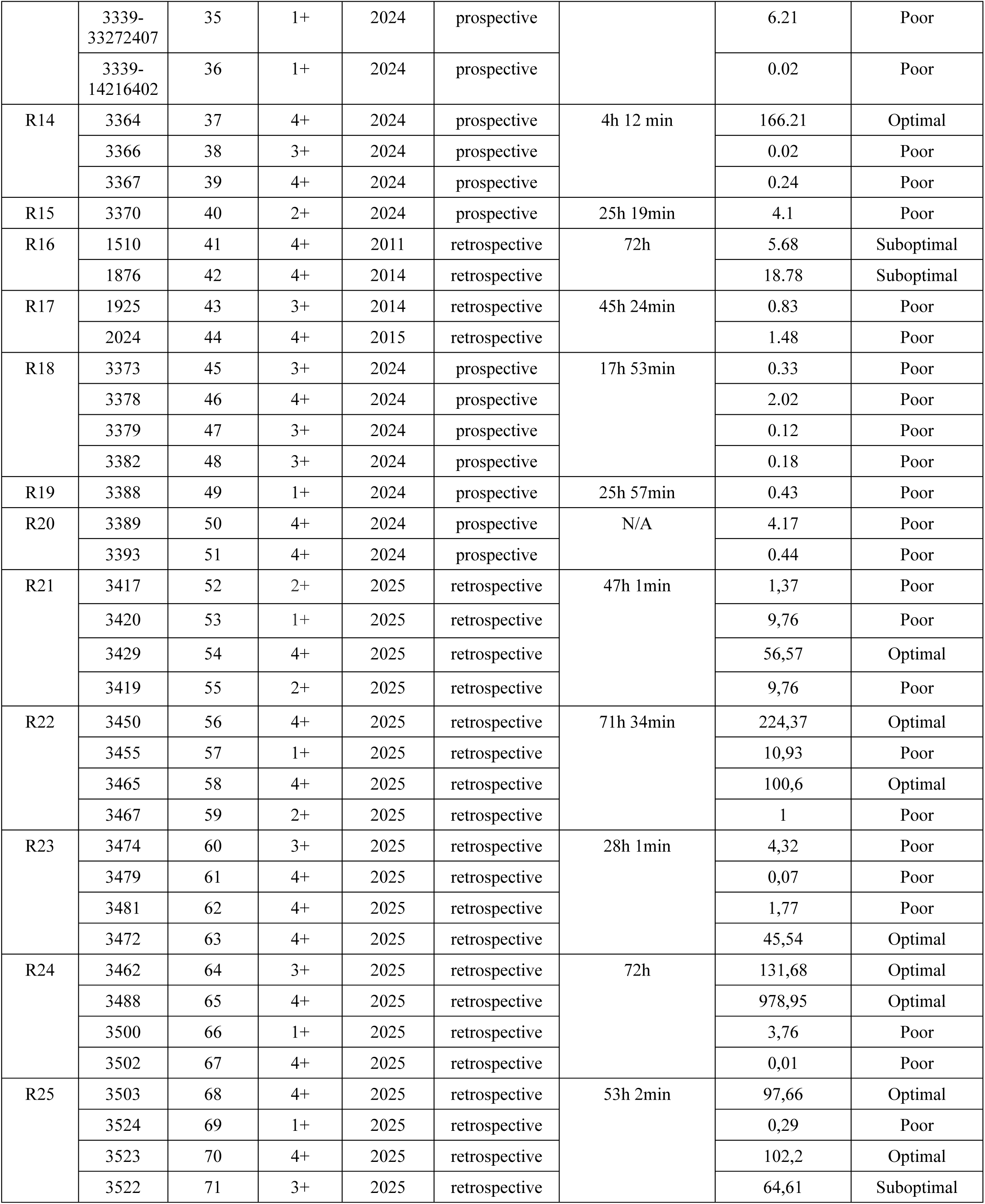
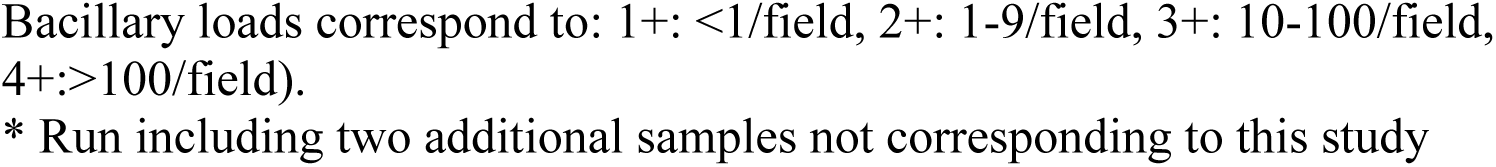
General performance of the 25 nanopore sequencing runs for the 71 sputa analyzed.

| Run number | Patient ID | Sample number | Bacillary load | Year of Collection (prospective/retrospective) |  | Run time | Mean coverage | Performance category |
| --- | --- | --- | --- | --- | --- | --- | --- | --- |
| R1 | 33490676-2 | 1 | 4+ | 2023 | prospective | 22h 13min | 2.15 | Poor |
|  | 3255 | 2 | 4+ | 2024 | prospective |  | 21.95 | Suboptimal |
|  | 3257 | 3 | 2+ | 2024 | prospective |  | 0.74 | Poor |
| R2* | 3268 | 4 | 4+ | 2024 | prospective | 2h 44min | 0.02 | Poor |
|  | 3262 | 5 | 4+ | 2024 | prospective |  | 0.03 | Poor |
|  | 3265 | 6 | 4+ | 2024 | prospective |  | 0.01 | Poor |
| R3 | 3281 | 7 | 4+ | 2024 | prospective | 26h 44min | 0.16 | Poor |
|  | 3290 | 8 | 3+ | 2024 | prospective |  | 1.04 | Poor |
| R4 | 1597 | 9 | 3+ | 2012 | retrospective | 3h 14min | 1.92 | Poor |
| R5 | 1165 | 10 | 4+ | 2009 | retrospective | 24h 27min | 825.29 | Optimal |
|  | 3278 | 11 | 4+ | 2024 | prospective |  | 29.2 | Suboptimal |
|  | 3280 | 12 | 4+ | 2024 | prospective |  | 3.88 | Poor |
| R6 | 691 | 13 | 3+ | 2007 | retrospective | 4h 17min | 69.9 | Optimal |
| R7 | 1598 | 14 | 4+ | 2012 | retrospective | 49h 20min | 10.7 | Suboptimal |
|  | 755 | 15 | 3+ | 2007 | retrospective |  | 15.53 | Suboptimal |
|  | 3299 | 16 | 4+ | 2024 | prospective |  | 489.2 | Optimal |
| R8 | 3313 | 17 | 4+ | 2024 | prospective | 24h 50min | 1.56 | Poor |
|  | 3315 | 18 | 4+ | 2024 | prospective |  | 5.76 | Suboptimal |
|  | 3296 | 19 | 4+ | 2024 | prospective |  | 816.69 | Optimal |
| R9 | 3283 | 20 | 4+ | 2024 | prospective | 28h 31min | 128.29 | Optimal |
|  | 3292 | 21 | 2+ | 2024 | prospective |  | 0.07 | Poor |
|  | 630 | 22 | 3+ | 2007 | retrospective |  | 0.7 | Poor |
|  | 3252 | 23 | 4+ | 2024 | prospective |  | 8.37 | Suboptimal |
| R10 | 3302 | 24 | 4+ | 2024 | prospective | 25h 10min | 144.05 | Optimal |
|  | 3319 | 25 | 4+ | 2024 | prospective |  | 1141.39 | Optimal |
|  | 3295 | 26 | 4+ | 2024 | prospective |  | 70.64 | Optimal |
| R11 | 3309 | 27 | 3+ | 2024 | prospective | 29h 22min | 99.05 | Optimal |
|  | 3325 | 28 | 4+ | 2024 | prospective |  | 122.07 | Optimal |
| R12 | 3324 | 29 | 2+ | 2024 | prospective | 72h | 3.46 | Poor |
|  | 3330 | 30 | 4+ | 2024 | prospective |  | 943.65 | Optimal |
|  | 3333 | 31 | 3+ | 2024 | prospective |  | 124.38 | Optimal |
|  | 3334 | 32 | 4+ | 2024 | prospective |  | 221.76 | Optimal |
| R13 | 3344 | 33 | 2+ | 2024 | prospective | 69h 59 min | 127.68 | Optimal |
|  | 3345 | 34 | 4+ | 2024 | prospective |  | 2863.31 | Optimal |
|  | 3339-33272407 | 35 | 1+ | 2024 | prospective |  | 6.21 | Poor |
|  | 3339-14216402 | 36 | 1+ | 2024 | prospective |  | 0.02 | Poor |
| R14 | 3364 | 37 | 4+ | 2024 | prospective | 4h 12 min | 166.21 | Optimal |
|  | 3366 | 38 | 3+ | 2024 | prospective |  | 0.02 | Poor |
|  | 3367 | 39 | 4+ | 2024 | prospective |  | 0.24 | Poor |
| R15 | 3370 | 40 | 2+ | 2024 | prospective | 25h 19min | 4.1 | Poor |
| R16 | 1510 | 41 | 4+ | 2011 | retrospective | 72h | 5.68 | Suboptimal |
|  | 1876 | 42 | 4+ | 2014 | retrospective |  | 18.78 | Suboptimal |
| R17 | 1925 | 43 | 3+ | 2014 | retrospective | 45h 24min | 0.83 | Poor |
|  | 2024 | 44 | 4+ | 2015 | retrospective |  | 1.48 | Poor |
| R18 | 3373 | 45 | 3+ | 2024 | prospective | 17h 53min | 0.33 | Poor |
|  | 3378 | 46 | 4+ | 2024 | prospective |  | 2.02 | Poor |
|  | 3379 | 47 | 3+ | 2024 | prospective |  | 0.12 | Poor |
|  | 3382 | 48 | 3+ | 2024 | prospective |  | 0.18 | Poor |
| R19 | 3388 | 49 | 1+ | 2024 | prospective | 25h 57min | 0.43 | Poor |
| R20 | 3389 | 50 | 4+ | 2024 | prospective | N/A | 4.17 | Poor |
|  | 3393 | 51 | 4+ | 2024 | prospective |  | 0.44 | Poor |
| R21 | 3417 | 52 | 2+ | 2025 | retrospective | 47h 1min | 1,37 | Poor |
|  | 3420 | 53 | 1+ | 2025 | retrospective |  | 9,76 | Poor |
|  | 3429 | 54 | 4+ | 2025 | retrospective |  | 56,57 | Optimal |
|  | 3419 | 55 | 2+ | 2025 | retrospective |  | 9,76 | Poor |
| R22 | 3450 | 56 | 4+ | 2025 | retrospective | 71h 34min | 224,37 | Optimal |
|  | 3455 | 57 | 1+ | 2025 | retrospective |  | 10,93 | Poor |
|  | 3465 | 58 | 4+ | 2025 | retrospective |  | 100,6 | Optimal |
|  | 3467 | 59 | 2+ | 2025 | retrospective |  | 1 | Poor |
| R23 | 3474 | 60 | 3+ | 2025 | retrospective | 28h 1min | 4,32 | Poor |
|  | 3479 | 61 | 4+ | 2025 | retrospective |  | 0,07 | Poor |
|  | 3481 | 62 | 4+ | 2025 | retrospective |  | 1,77 | Poor |
|  | 3472 | 63 | 4+ | 2025 | retrospective |  | 45,54 | Optimal |
| R24 | 3462 | 64 | 3+ | 2025 | retrospective | 72h | 131,68 | Optimal |
|  | 3488 | 65 | 4+ | 2025 | retrospective |  | 978,95 | Optimal |
|  | 3500 | 66 | 1+ | 2025 | retrospective |  | 3,76 | Poor |
|  | 3502 | 67 | 4+ | 2025 | retrospective |  | 0,01 | Poor |
| R25 | 3503 | 68 | 4+ | 2025 | retrospective | 53h 2min | 97,66 | Optimal |
|  | 3524 | 69 | 1+ | 2025 | retrospective |  | 0,29 | Poor |
|  | 3523 | 70 | 4+ | 2025 | retrospective |  | 102,2 | Optimal |
|  | 3522 | 71 | 3+ | 2025 | retrospective |  | 64,61 | Suboptimal |

Bacillary loads correspond to: 1+: <1/field, 2+: 1-9/field, 3+: 10-100/field, 4+:>100/field). \* Run including two additional samples not corresponding to this study

Among the study samples, 24 of them (33.8%) led to optimal results (>90% of genome covered, mean coverage >45 × and >70% genome covered >20× ; Figure 1); the shortest run that allowed us to achieve >90% at mean coverage depth >50 × in a sample took 4 hours and 12 minutes (Table 1). Another 9 samples (12.6%) yielded suboptimal results, with a range of 15.5% to 90.92% genome covered at >10 ×. The remaining 38 sputa (53.5%) were classified as poor results, with genome covered at ≤ 15% at >10 × (see Supplementary table).

**Figure 1:**
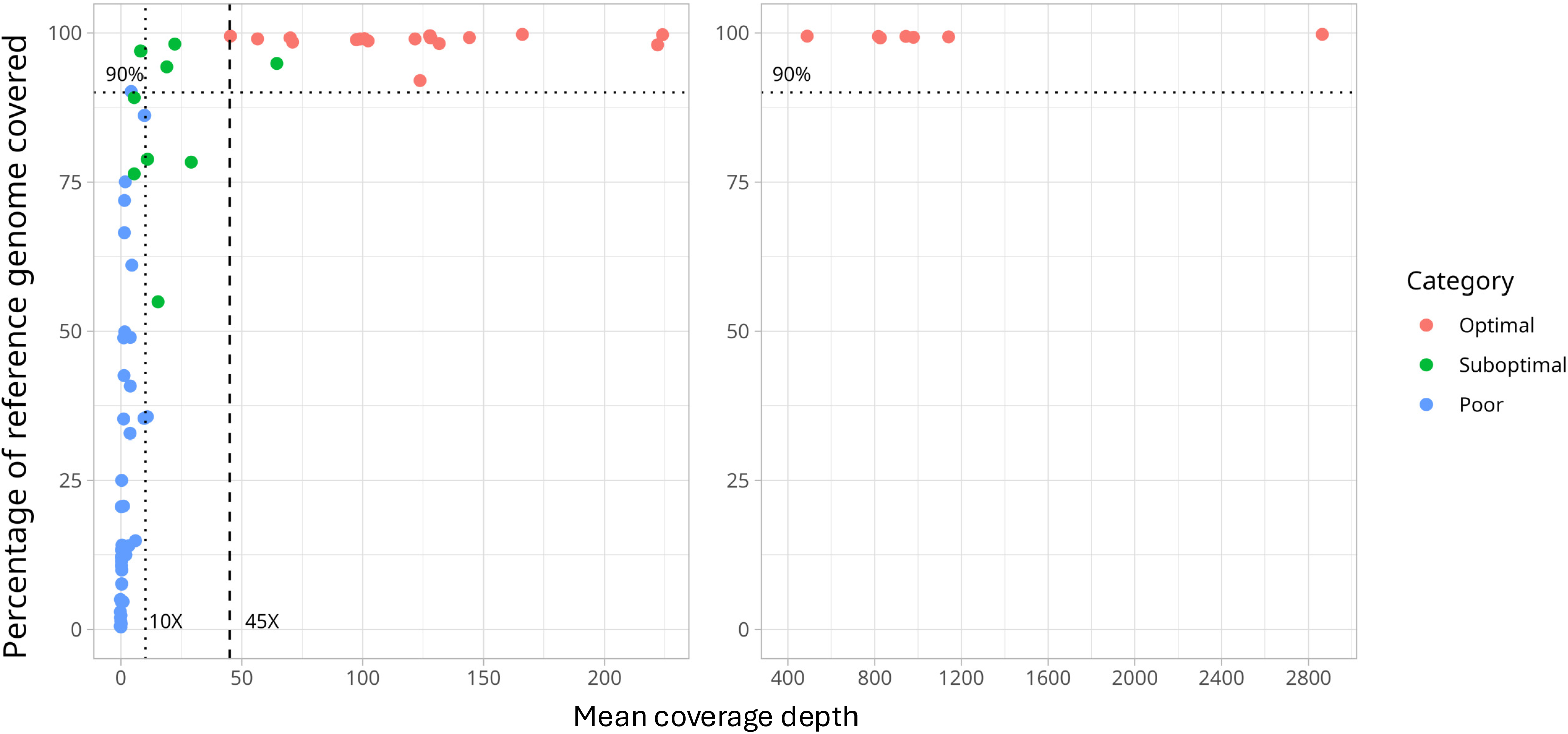
Sequencing performance (percentage of the MTB reference genome that is covered and mean depth of coverage), for the specimens in the study that have been categorised as optimal, suboptimal, or poor. Each dot in the graph corresponds to one sputum sample.

For each category (optimal, suboptimal and poor), the percentage of reads corresponding to human, bacteria other than MTB, and MTB was determined (Figure 2). Despite incorporating a procedure that depletes human DNA in the extraction stage, a substantial variation in efficiency was observed for the depletion. Optimal results were obtained even in samples containing a majority proportion of interfering DNA from either other bacteria or human. The preponderance of poor outcomes was observed to be associated with samples exhibiting a low relative proportion of MTB DNA.

**Figure 2:**
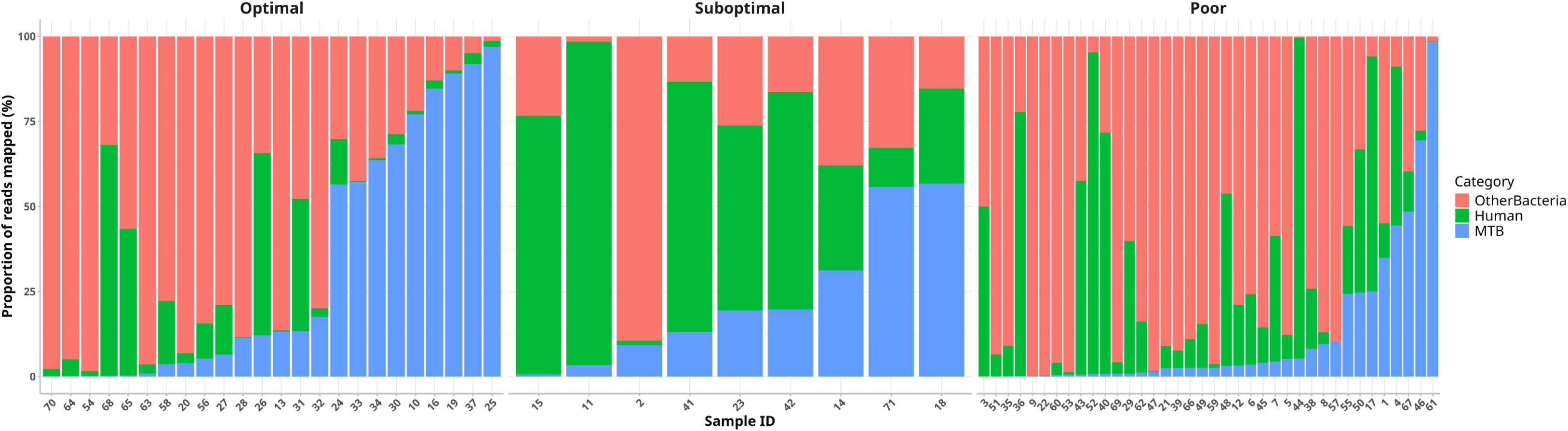
Proportion of reads mapping with MTBC, human, or bacteria from the sequences obtained in each sputum sample for the optimal, suboptimal, and poor sequencing categories.

### Evaluation of factors that might predict sequencing performance

Our aim was to determine whether certain factors could be utilised to predict when sequencing from sputa is likely to yield optimal, suboptimal, or poor results. With regard to the bacillary load, as determined by smear grade, the majority of the samples (76%) were classified as 3+/4+ specimens (Table 1). This precluded an assessment of the potential association between bacterial load and sequencing performance. However, it was observed that in all but one of the cases with low bacillary load, poor results were obtained, while all the optimal results corresponded to higher bacillary loads (Table 1). It is also true that poor results were obtained from sputa with high bacillary loads, indicating that smear grade by itself does not predict the quality of the results. Therefore, we aimed to estimate, in addition to the presence of MTB in the samples, the presence of interfering DNA. This was achieved by examining the Ct values for three real-time PCRs targeting bacteria, human DNA and MTB (Figure 3).

**Figure 3:**
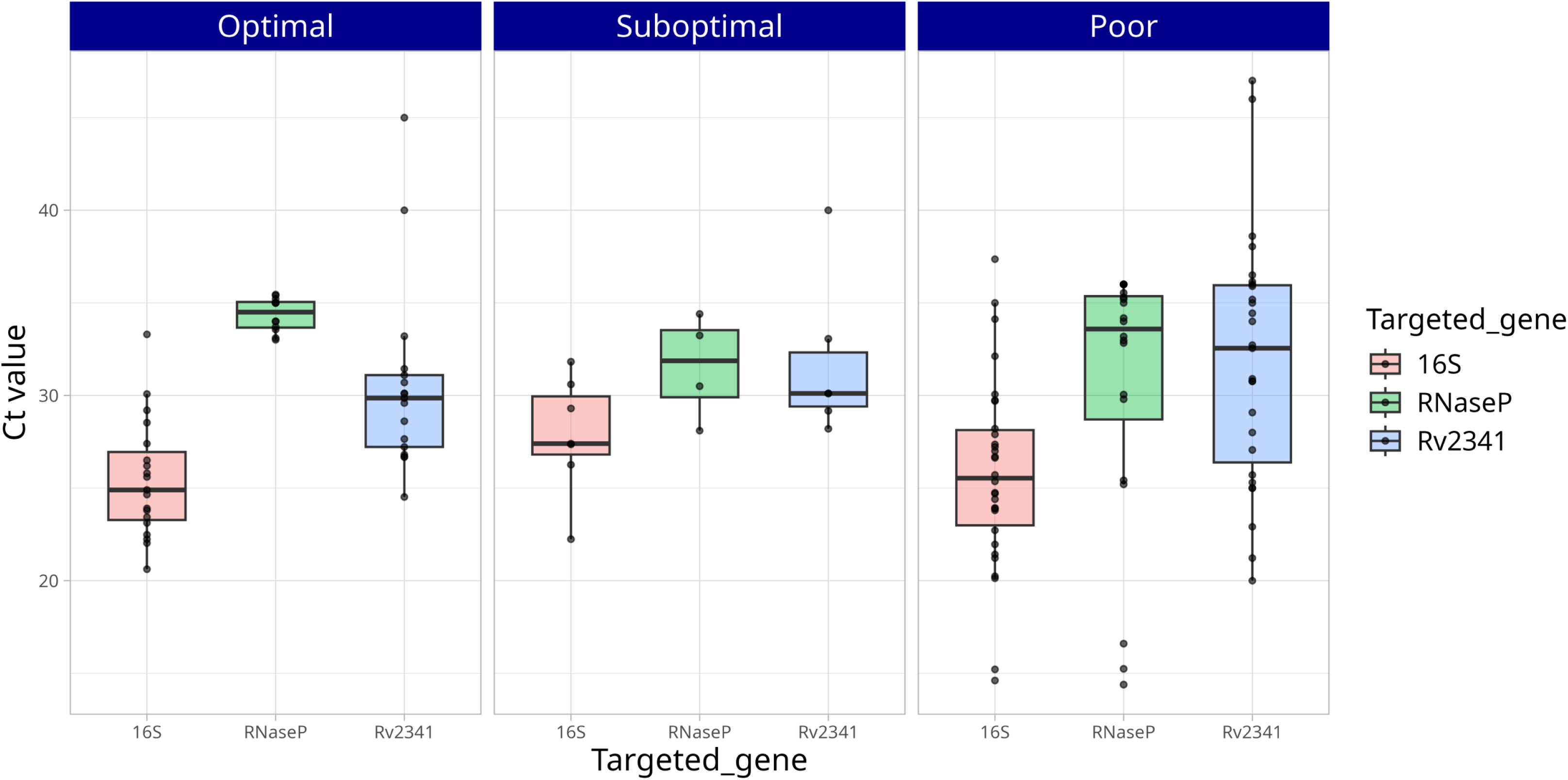
Ct values for RT-PCRs targeting 16S rRNA, RNase P, and Rv2341 for the optimal, suboptimal, and poor sequencing categories.

As was previously observed, the efficiency of the human depletion steps was found to be variable, resulting in significant variation in the *RNase P* Ct values. The distribution of the Ct values for human DNA was analysed, and it was observed that optimal results were obtained when there was a lower presence of human DNA. However, the distribution of the relative proportions of DNA from either MTBC or bacteria was equivalent between the specimens, leading to optimal or suboptimal results. It was also found that the distribution was especially diverse in those corresponding to poor results. In summary, equivalent Ct values for MTBC may lead to optimal or poor results, depending on the presence of either accompanying bacteria or human material. Poor results were those in which we observed the highest representativity of human/bacterial DNA together with the lowest proportions of MTBC DNA. These observations, altogether, indicate that it is challenging to find a straightforward way to predict the sequencing performance expected for a specimen.

### Allocating cases to clusters

#### Assignment from optimal sequences

In the 24 cases that demonstrated optimal results, whether from the prospective or retrospective rounds of analysis we could determine that six of these cases (cases 3283, 3295, 3325, 3472, 3330 and 3465) corresponded to clustered cases (Clusters 3133, 899-2204, 3325, 3330 and 1414), while the remaining cases were designated as orphan cases.

#### Rescue of data from suboptimal sequences

It is acknowledged that complete genome analysis is not feasible when suboptimal results are obtained. We aimed to evaluate the potential for partial exploitation of suboptimal, at least to facilitate the pre-assignment of cases to existing clusters. The rationale behind this approach was to attempt to identify SNPs from suboptimal sequences that are markers for preexisting clusters.

In order to address this purpose, the more epidemiologically relevant clusters in the population were first selected, namely those which were either newly identified or had grown (incorporating ≥1 case, 1-7 cases) over the last five years (2021-2025). A total of 34 clusters were selected (Table 2).

**Table 2:**
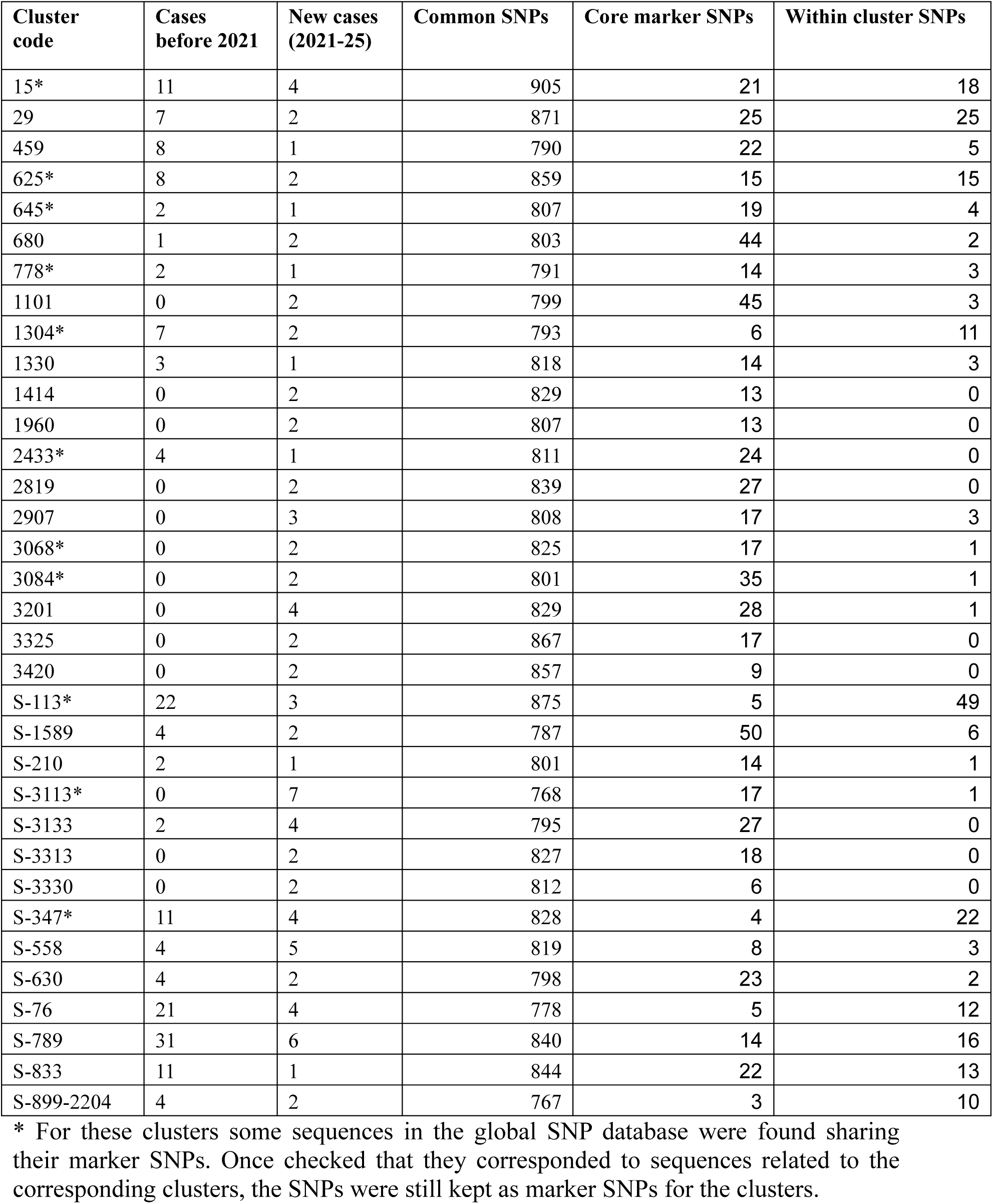
Common, core marker and within cluster SNPs for the epidemiologically relevant clusters selected.

| Cluster code | Cases before 2021 | New cases (2021-25) | Common SNPs | Core marker SNPs | Within cluster SNPs |
| --- | --- | --- | --- | --- | --- |
| 15* | 11 | 4 | 905 | 21 | 18 |
| 29 | 7 | 2 | 871 | 25 | 25 |
| 459 | 8 | 1 | 790 | 22 | 5 |
| 625* | 8 | 2 | 859 | 15 | 15 |
| 645* | 2 | 1 | 807 | 19 | 4 |
| 680 | 1 | 2 | 803 | 44 | 2 |
| 778* | 2 | 1 | 791 | 14 | 3 |
| 1101 | 0 | 2 | 799 | 45 | 3 |
| 1304* | 7 | 2 | 793 | 6 | 11 |
| 1330 | 3 | 1 | 818 | 14 | 3 |
| 1414 | 0 | 2 | 829 | 13 | 0 |
| 1960 | 0 | 2 | 807 | 13 | 0 |
| 2433* | 4 | 1 | 811 | 24 | 0 |
| 2819 | 0 | 2 | 839 | 27 | 0 |
| 2907 | 0 | 3 | 808 | 17 | 3 |
| 3068* | 0 | 2 | 825 | 17 | 1 |
| 3084* | 0 | 2 | 801 | 35 | 1 |
| 3201 | 0 | 4 | 829 | 28 | 1 |
| 3325 | 0 | 2 | 867 | 17 | 0 |
| 3420 | 0 | 2 | 857 | 9 | 0 |
| S-113* | 22 | 3 | 875 | 5 | 49 |
| S-1589 | 4 | 2 | 787 | 50 | 6 |
| S-210 | 2 | 1 | 801 | 14 | 1 |
| S-3113* | 0 | 7 | 768 | 17 | 1 |
| S-3133 | 2 | 4 | 795 | 27 | 0 |
| S-3313 | 0 | 2 | 827 | 18 | 0 |
| S-3330 | 0 | 2 | 812 | 6 | 0 |
| S-347* | 11 | 4 | 828 | 4 | 22 |
| S-558 | 4 | 5 | 819 | 8 | 3 |
| S-630 | 4 | 2 | 798 | 23 | 2 |
| S-76 | 21 | 4 | 778 | 5 | 12 |
| S-789 | 31 | 6 | 840 | 14 | 16 |
| S-833 | 11 | 1 | 844 | 22 | 13 |
| S-899-2204 | 4 | 2 | 767 | 3 | 10 |

\* For these clusters some sequences in the global SNP database were found sharing their marker SNPs. Once checked that they corresponded to sequences related to the corresponding clusters, the SNPs were still kept as marker SNPs for the clusters.

Subsequently, the marker SNPs for the relevant clusters were identified, utilising the genomic sequences obtained before, from the cases from the selected clusters, within our genomic surveillance programme (17). Firstly, the common SNPs shared by all members in each cluster were determined (Table 2). Then, these were compared with an in-house database containing the SNPs from 109,648 sequences taken from publicly available databases, representing 121 countries from the year 1909 to the year 2024. Following this comparison, all common cluster common SNPs found in the database were filtered out, to obtain a list of a total of 641 key unique SNPs (3-50 core marker SNPs/cluster, only found in the members of each cluster; Table 2). Furthermore, the list was expanded to encompass the differential SNPs identified between the cases included in each of the clusters (230 within-cluster SNPs). The total number of informative SNPs compiled for the 34 clusters selected was 871 (Table 2).

Based on these results, a marker SNP pipeline (rescue pipeline) was customised for the purpose of calling the informative cluster SNPs from the nine suboptimal sequences that were obtained in the course of the analysis. From five of the nine cases exhibiting suboptimal sequences (1598, 755, 3315 and 3252, and 3522, Table 3), a proportion of the core marker SNPs could be called, while no calls were obtained for the remaining four cases. Among the five samples with calls, in four of them, sufficient coverage (at ≥ 5×) was obtained to call 33, 2, 7 and 23 SNPs, respectively, of the corresponding cluster core marker SNPs (30%, 40%, 87.5% and 91.3% of their total cluster marker SNPs; Table 3). These calls facilitated the allocation of these subjects as probable candidates for entry into four preexisting active clusters (Clusters 1589, 113, 558, and 630). Furthermore, for patient 3252, who had been preassigned to Cluster 630 by the rescue pipeline, it also enabled us to allocate it within the corresponding genomic network (Figure 4). The correct call from the suboptimal sequences from patient 3252 of all the SNPs called in Cases 630 and 912, in conjunction with one new SNP detected as acquired after analysing the suboptimal sequences indicated that Case 3252 was the last case in the network (Figure 4). For the remaining case (case 3522), only 20% of the marker SNPs from one of the relevant clusters were called at > 5x, (Cluster 29; Table 3), which was not considered sufficient for a pre-assignation.

**Figure 4:**
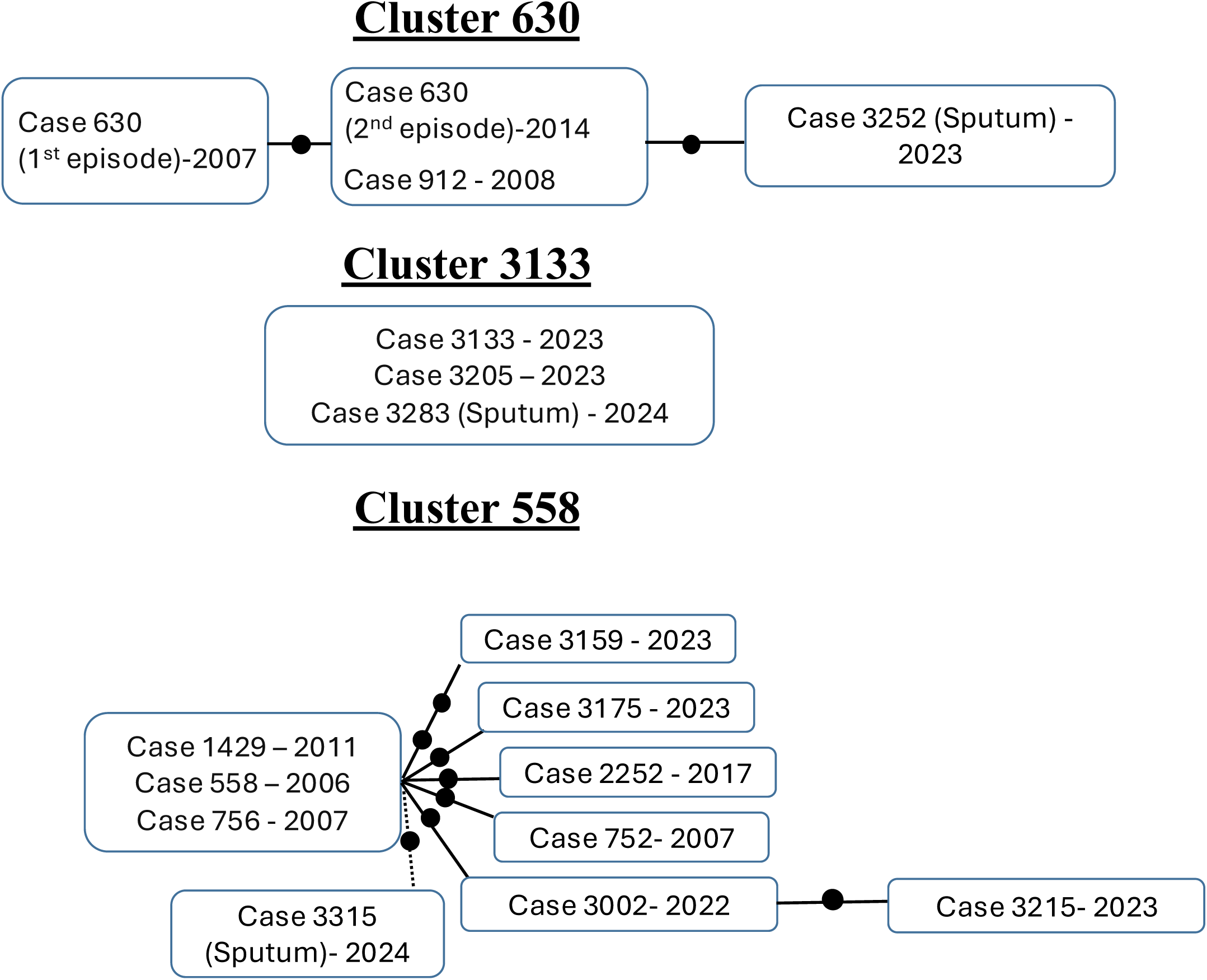
Genomic networks for the pre-existing clusters in which a new case was identified in the prospective sub-study (each dot corresponds to 1 SNP; the cases within the same box had 0 SNPs among them). Cases assigned to these clusters from sequencing analysis on sputa are indicated (sputum). In cluster 558, suboptimal sequences from Case 3315 allowed us to assign it to this cluster, but the differential SNP drawn (dotted line) was obtained from the subsequent sequencing on culture.

**Table 3:** Samples leading to suboptimal sequences that allowed us to call Cluster informative SNPs by applying the rescue pipeline.

| Sample Number | Patient ID | Cluster | Core marker SNPs | Within cluster marker SNPs | Total called_SNPs | Called_Core_SNPs %≥5X | Total SNPs ≥ 1X | Total SNPs ≥ 5X | Total SNPs ≥10X |
| --- | --- | --- | --- | --- | --- | --- | --- | --- | --- |
| 14 | 1598 | 1589 | 50 | 6 | 33 core | 30,00 % | 33 | 15 | 8 |
| 15 | 755 | 113 | 5 | 49 | 2 core | 40,00 % | 2 | 2 | 2 |
| 18 | 3315 | 558 | 8 | 4 | 7 core | 87,50 % | 7 | 4 | 1 |
| 23 | 3252 | 630 | 23 | 2 | 23 core + 1 within cluster | 91,30 % | 23 | 21 | 11 |
| 71 | 3522 | 29 | 25 | 25 | 6 core | 20,00 % | 6 | 5 | 0 |

In all cases, including this last case 3522, in which a decision was taken not to pre-assign it to Cluster 29, these preliminary assignments made by the rescue pipeline were subsequently confirmed from the WGS data, once the corresponding cultures were available.

#### Epidemiological exploitation of results from sputa

The samples included in the prospective subanalysis presented in this study enabled the evaluation of the accelerated allocation of new cases into three preexisting clusters (cases 3252, 3283 and 3315 in clusters 630, 3133 and 558, respectively).

Cluster 630 (Figure 4) involved two preceding migrant cases (cases 630 and 912), two siblings with TB in 2007 and 2008, respectively. Case 630 exhibited a second TB episode in 2014, consequent to inadequate adherence to anti-TB treatment during the initial episode. The inclusion of case 3252 in this cluster prompted a rapid evaluation of its relationship with the preceding cases. Indeed, as a consequence of this finding, Case 3252 was interviewed, and it was revealed that he had been a close contact of Case 630 during the latter’s second episode in 2014. However, he did not adhere to prophylactic treatment at that time. The rapid genomic analysis of sputum samples enabled the diagnosis of this case as a reactivation resulting from a prior exposure and, therefore, not wasting the specific control efforts that are dedicated when recent active transmissions are identified, to identify new non-diagnosed active cases.

Cluster 3133 (Figure 4) corresponded to a recent active transmission event involving two recently arrived migrant cases (cases 3133 and 3205, diagnosed in 2023 and 2024, 1.5 and 5 years after arrival, respectively) residing in a context characterised by a high proportion of migrant population. The two preceding cases exhibited 0 SNPs between them; the inclusion of case 3283 in this cluster (also exhibiting 0 SNPs in relation to the preceding cases) indicated his involvement in the same recent common transmission event. In the absence of the rapid genomic assignment of Case 3283 to this cluster, his TB would have been interpreted as a reactivation of a past exposure. This is because Case 3283 had resided in Almería for 22 years prior to his diagnosis, and a type II diabetes diagnosis was made in 2024, coinciding with his diagnosis of TB.

Cluster 558 was an extensive cluster involving 9 preceding cases (mainly migrants). The star-like topology of this cluster network (Figure 4) is consistent with sequential reactivations, for most of the cases, several years after exposure to an active transmission event, that happed years before (during 2006-07). The fast allocation of case 3315 to this cluster based on genomic analysis in sputum, prompted consideration of reactivation also as the most probable explanation for this case, a phenomenon that occurred in all but one of the anteceding cases in this cluster. This hypothesis was consistent with the finding of a poorly controlled diabetes and the patient’s advanced age (60 years). Our preliminary consideration of reactivation based on the genomic data obtained from sputa were finally reinforced, once the sequencing data from the corresponding culture were later available. These final consolidated data allowed us to locate Case 3315 in the network (Figure) directly linked with the cases involved in the active transmission event which occured in the past, also following the previously mentioned star-like topology.

## Discussion

Once WGS demonstrated its following the demonstration of the value of WGS in the identification of resistance mutations in MTB and the tracking of TB transmission with the utmost precision, efforts have been made to accelerate the availability of genomic data. The first movement was to conduct the analysis directly on primary cultures (18), obviating the necessity for subculturing. In an effort to circumvent the high-throughput schemes that cause unavoidable delays due to the necessity to accumulate a substantial number of cultures to be executed concurrently, a more flexible nanopore sequencing approach has been proposed (19). However, in order to reduce the time lag between new diagnoses and the availability of genomic data, it is necessary to adopt a culture-free strategy, by sequencing directly on sputa (20). It is evident that this would provide the most expeditious outcomes, whilst concomitantly circumventing constraints in settings lacking access to biosafety facilities and, consequently, impeding the implementation of culture-based diagnostics.

The process of direct sequencing of MTB in sputa is a challenging task due to the presence of high amounts of accompanying DNA from either human cells or other bacteria in the respiratory specimens. This accompanying DNA can interfere with the sequencing process. In the first study, which established the feasibility of directly sequencing MTB from clinical specimens, 20-93% of the obtained sequences corresponded to human reads (21), despite the implementation of a procedure aimed at depleting human DNA present in the specimen. This finding elucidated the remarkably low coverage obtained for MTB, which even fell below the mean coverage threshold of 0.7 × .

These suboptimal outcomes led to the conclusion that, when attempting to sequence directly on sputa, the enrichment in MTB provided by culturing should be substituted by alternative enrichment procedures (20). The aforementioned enrichments have relied on indirect approaches, as in the previously mentioned article, by eliminating the interfering human DNA through differential lysis procedures (18, 21, 22), or on direct schemes, by selectively capturing MTB by ligand-magnetic binding beads (23), or by hybridising with specifically designed capture probes (biotinylated RNA baits) to selectively capture MTB libraries before sequencing (22, 24, 25).

The application of an optimised procedure for the depletion of human DNA (18) resulted in enhanced coverage when conducting direct sequencing of sputa. However, the most favourable outcomes in terms of coverage depth and breadth have been achieved using RNA baits (22, 24, 25), despite the fact that its application is linked to lengthier, more cumbersome and costly procedures.

To date, the scarcity of research in this field is evidenced by the paucity of studies addressing the challenge of directly sequencing MTB in sputa. This situation is particularly pertinent in low- and medium-burden settings, where further analysis is required. Moreover, the preponderance of this research has been oriented towards the evaluation of the efficiency with which resistance mutations can be identified, with a number of studies also incorporating preliminary analyses of lineage assignment or the construction of phylogenies (23–26). However, the evaluation of culture-free sequencing in the context of genomic epidemiology has received minimal attention. It is evident that the acceleration of precise, personalised therapy for each patient has a discernible impact on their prognosis, thereby validating the endeavours to expedite culture-free analysis. However, in contexts such as Almeria, where epidemiological interventions are coupled and oriented by the genomic characterisation of every new case (17), it is equally relevant to accelerate the genomic analysis. The timely identification and analysis of new cases is therefore paramount to orientate accordingly the effectiveness of contact tracing. This enables the enhancement of the identification and treatment of LTI cases, which in turn reduces the number of secondary cases and the overall burden of the disease. However, to the best of our knowledge, only one study (22) has addressed the culture-free genomic inference of transmission; the authors followed an innovative rationale, restricting the application of capturing-baits for the sputa with a low representativity of MTB DNA and following a direct non-capture approach for the remaining. In both branches, Illumina sequencing was employed.

The objective of our study is threefold: firstly, to address the paucity of studies dealing with culture-free sequencing of MTB in the context of genomic epidemiology; secondly, to accelerate the inference of new cases entering transmission clusters; and thirdly, to enhance our interventionist dynamic as soon as a new clustered case is identified. Looking for this acceleration, our present study proposes an alternative approach to the conventional Illumina-based sequencing method, namely nanopore sequencing, which has the potential to enhance the analysis’s efficiency and expedite the process. A mere one of the studies on culture-free MTB sequencing is supported on nanopore sequencing (26), and it only provided rather suboptimal results because the authors decided against applying any enrichment method, in an attempt to adapt to the limited resources found in many high-burden settings. The hypothesis that nanopore is better adapted to a faster response has been demonstrated by our team in recent studies using primary cultures. This finding suggests that a more flexible case-by-case analysis of new incident cases can be performed (19). Votintseva *et al.* also established that nanopore provided a faster response, although this finding was only supported by a pilot study using BCG spiked sputa (18).

Consequently, the present study constitutes the first analysis of culture-free sequencing of MTB with a genomic epidemiology scope supported by flexible nanopore sequencing. In this study, a depletion approach was employed; instead of a capture enrichment method, with the objective of offering a less complex and less expensive procedure. The number of specimens evaluated, 71, was the highest among other previous studies. In one-third of the cases, the quality of our results was equivalent to that obtained when sequencing directly in cultures, thus enabling us to perform a complete comparative genomics analysis. It is noteworthy that we were able to achieve substantial coverage from a 2+ specimen, and in some instances, good coverages were accomplished with short run-times. It is important to acknowledge that in approximately half of the cases under consideration, no exploitable results, for our comparative genomics purposes, were obtained. However, it should be noted that the coverage values obtained in other studies were even below the less demanding thresholds required for tasks that are much more straightforward than comparative genomics, such as the identification of resistance mutations or the assignment of lineages (26). Furthermore, even when employing sophisticated techniques, such as the use of magnetic beads coupled with ligands to bind and enrich MTB (23), the maximum coverages attained were only 18.7× and the overall genome coverage was 15.2%, thereby hindering even straightforward tasks, such as the assignment of lineages, which was only possible in 15% of the samples. Even some of the studies among those achieving the best results, by enriching MTB libraries by using RNA-baits, faced room for improvement, with still 24% of the samples not reaching the thresholds for the purpose of comparative genomics (22).

We aimed to ascertain the underlying reasons for our proportion of unsuccessful results. Firstly, by evaluating the performance depending on the bacillary load, as inferred by the staining. Optimal results were only obtained in one sputum sample with a low bacillary load. All subsequent optimal results were obtained from sputum samples with higher bacillary loads. However, the performance of the sputa with higher loads ranged from poor to optimal, thus impairing predictions. Subsequently, the distribution of reads obtained from human/other accompanying bacteria was determined, as well as the relative proportion of MTB, bacterial and human DNA in the extracted samples. Our initial observation revealed that, despite the implementation of a differential lysis-based kit to deplete human DNA, the kit’s performance exhibited significant variability. This variability was characterised by the presence of high percentages of human DNA, or low Ct values, among the reads obtained from certain samples. The presence of human DNA even after the application of depletion procedures has also been reported in other studies (21), indicating the need for methodological improvements at this stage. The variable presence of human and bacterial DNA hindered the establishment of a criterion to predict whether a sputum sample would yield optimal or poor sequencing results. A similar absence of correlation between bacterial load or smear grade and sequencing coverage has been identified in other studies (26). A correlation could be identified in studies that employed enrichment by RNA baits (23), thereby enabling the authors to minimise the interference of human DNA and consequently ascertain the correlation.

It is acknowledged that obtaining WGS data directly from sputa is currently challenging (26) and that a proportion of culture-free results are expected to be suboptimal. In light of these issues, an innovative rescue pipeline was proposed by us to exploit these sequences not reaching the thresholds required for comparative genomics. The rationale behind the rescue pipeline is to identify informative SNPs in genomic regions that are adequately covered in the sequences, despite suboptimal average coverage values. The SNPs deemed informative are those which were identified to be exclusively present in the relevant clusters within our population; consequently, they can function as cluster-marker-SNPs. Consequently, if we can call them in a set of suboptimal sequences, we can preassign the corresponding case to a preexisting relevant cluster. The potential of marker SNPs to preassign a strain to a certain cluster has been demonstrated by our team in many preceding studies and settings. This has been achieved by using targeted PCRs to track relevant strains in France, Panama, Costa Rica, Morocco, Belgium, Peru, and others in a faster, simpler, and low-cost way (17, 27–31). Marker SNPs have also been proposed as a means of facilitating more efficient and expeditious identification of the primary MDR strains circulating in Europe (32). The current rescue pipeline utilises the same concepts but now seeks these marker SNPs among the calls obtained from samples with suboptimal results. This approach enabled us to identify cases of preexisting clusters in several instances, and in one particular instance, it facilitated the precise location of the case within the genomic network of the cluster. In all these cases, the pre-assignments, supported by the identification of a subset of marker SNPs, were validated by the complete sequencing data obtained at a later stage, once the cultures were available.

It is true that the inference of clusters in a population from culture-free data, particularly in the context of implementing a rescue pipeline, necessitates the pre-existence of a long-term surveillance programme. This programme facilitates the identification of the most relevant clusters, as determined by their size and the rate at which new cases have entered those clusters. In this particular instance, the rapid identification of genomic assignments for new cases to preexisting clusters, within the prospective subanalysis, offered special value. Indeed, the findings of this study have enabled us to ascertain when the new entrances were more likely to be the result of either reactivations due to past exposures or active recent transmission. This recommended that special efforts to enhance transmission control should particular focus on the cluster associated with active transmission. The new entrances in the other two clusters, attributable to the reactivation of prior exposures, underscores the challenges inherent in ensuring comprehensive coverage in contact tracing within these epidemiologically intricate populations characterised by high rates of migration, and consequently, to ensuring the administration of prophylactic treatment for tuberculosis infection in all pertinent cases.

In summary, the present study provides additional insights into the possibilities of culture-free whole genome sequencing of *Mycobacterium tuberculosis* in the context of interventionist precision genomic epidemiology. Despite the identification of scope for enhancement, the findings of this study initiate the process of integrating a rapid analysis in close proximity to diagnosis and a preliminary categorisation of cases, despite suboptimal outcomes.

## Acknowledgements

CIBER - Consorcio Centro de Investigación Biomédica en Red (CB06/06/0058), COST-Action-AdvanceTB (CA21164), and Computing facilities at CETA-CIEMAT with ERDF funds.

## Data availability statement

The sequences generated for Nanopore were deposited in the ENA (project number PRJEB97396). Human reads have been removed from the sequences to maintain anonymity of the sequence.

## References

1. Jajou R, De Neeling A, Van Hunen R, De Vries G, Schimmel H, Mulder A, et al. Epidemiological links between tuberculosis cases identified twice as efficiently by whole genome sequencing than conventional molecular typing: A population-based study. PLoS ONE. 2018;13(4).

2. Doyle RM, Burgess C, Williams R, Gorton R, Booth H, Brown J, et al. Direct whole-genome sequencing of sputum accurately identifies drug-resistant mycobacterium tuberculosis faster than MGIT culture sequencing. J Clin Microbiol. 2018;56(8).

3. Vasanthaiah S, Verma R, Kumar A, Bandari AK, George J, Rastogi M, et al. Culture-Free Whole Genome Sequencing of Mycobacterium tuberculosis Using Ligand-Mediated Bead Enrichment Method. Open Forum Infect Dis. 2024 Jul 1;11(7).

4. Albert H, Ademun PJ, Lukyamuzi G, Nyesiga B, Manabe Y, Joloba M, et al. Feasibility of magnetic bead technology for concentration of mycobacteria in sputum prior to fluorescence microscopy. BMC Infect Dis. 2011;11.

5. Brown AC, Bryant JM, Einer-Jensen K, Holdstock J, Houniet DT, Chan JZM, et al. Rapid whole-genome sequencing of mycobacterium tuberculosis isolates directly from clinical samples. J Clin Microbiol. 2015;53(7).

6. Doughty EL, Sergeant MJ, Adetifa I, Antonio M, Pallen MJ. Culture-independent detection and characterisation of Mycobacterium tuberculosis and M. africanum in sputum samples using shotgun metagenomics on a benchtop sequencer. PeerJ. 2014;2014(1).

7. Nilgiriwala K, Rabodoarivelo MS, Hall MB, Patel G, Mandal A, Mishra S, et al. Genomic Sequencing from Sputum for Tuberculosis Disease Diagnosis, Lineage Determination, and Drug Susceptibility Prediction. J Clin Microbiol. 2023;61(3).

8. Bagratee TJ, Studholme DJ. Targeted genome sequencing for tuberculosis drug susceptibility testing in South Africa: a proposed diagnostic pipeline. Access Microbiol. 2024;6(2).

9. Cabibbe AM, Moghaddasi K, Batignani V, Morgan GSK, Di Marco F, Cirillo DM. Nanopore-based targeted sequencing test for direct tuberculosis identification,genotyping, and detection of drug resistance mutations: a side-by-side comparison of targeted next-generation sequencing technologies. Journal of Clinical Microbiology. 2024 Oct 1;62(10).

10. Votintseva AA, Bradley P, Pankhurst L, Del Ojo Elias C, Loose M, Nilgiriwala K, et al. Same-day diagnostic and surveillance data for tuberculosis via whole-genome sequencing of direct respiratory samples. J Clin Microbiol. 2017;55(5).

11. Mariner-Llicer C, Goig GA, Torres-Puente M, Vashakidze S, Villamayor LM, Saavedra-Cervera B, et al. Genetic diversity within diagnostic sputum samples is mirrored in the culture of Mycobacterium tuberculosis across different settings. Nature Communications. 2024 Dec 1;15(1).

12. Abascal E, Pérez-Lago L, Martínez-Lirola M, Chiner-Oms Á, Herranz M, Chaoui I, et al. Whole genome sequencing-based analysis of tuberculosis (TB) in migrants: Rapid tools for crossborder surveillance and to distinguish between recent transmission in the host country and new importations. Eurosurveillance. 2019;24(4).

13. Goig GA, Torres-Puente M, Mariner-Llicer C, Villamayor LM, Chiner-Oms Á, Gil-Brusola A, et al. Towards next-generation diagnostics for tuberculosis: Identification of novel molecular targets by large-scale comparative genomics. Bioinformatics. 2020;36(4).

14. Buenestado-Serrano S, Vallejo-Godoy S, Escabias Machuca F, Barroso P, Martínez-Lirola M, Cabezas T, et al. Redefinition of Transmission Clusters by Accessing to Additional Diversity in Mycobacterium Tuberculosis Through Long-Read Sequencing. 2025 [cited 2025 Aug 27]; Available from: https://papers.ssrn.com/abstract=5166258

15. Comas Ĩ, Chakravartti J, Small PM, Galagan J, Niemann S, Kremer K, et al. Human T cell epitopes of Mycobacterium tuberculosis are evolutionarily hyperconserved. Nature Genetics. 2010;42(6).

16. Leigh JW, Bryant D. POPART: Full-feature software for haplotype network construction. Methods in Ecology and Evolution. 2015;6(9).

17. Rodriguez Grande C, Vallejo Godoy S, Martinez Lirola M, Saleeb S, Herranz M, Buenestado Serrano S, et al. Long-term refined genomic analysis of tuberculosis clusters to distinguish between ongoing transmission, reactivations or diagnostic delays. Eurosurveillance. 2025;In press.

18. Votintseva AA, Bradley P, Pankhurst L, Del Ojo Elias C, Loose M, Nilgiriwala K, et al. Same-day diagnostic and surveillance data for tuberculosis via whole-genome sequencing of direct respiratory samples. Journal of Clinical Microbiology. 2017;55(5).

19. Rodríguez-Grande, Cristina, M. Saleeb, Sheri, Sanz-Pérez A, Buenestado-Serrano S, Martínez-Lirola M, Herranz-Martín M, Peñas-Utrilla D, Muñoz P, et al. Reducing delays in the genomic epidemiology of tuberculosis: a flexible and decentralized analysis of each incident case. Unpublished. 2024 Jul 19;

20. McNerney R, Clark TG, Campino S, Rodrigues C, Dolinger D, Smith L, et al. Removing the bottleneck in whole genome sequencing of Mycobacterium tuberculosis for rapid drug resistance analysis: a call to action. International journal of infectious diseases : IJID : official publication of the International Society for Infectious Diseases. 2017 Mar 1;56:130–5.

21. Doughty EL, Sergeant MJ, Adetifa I, Antonio M, Pallen MJ. Culture-independent detection and characterisation of Mycobacterium tuberculosis and M. africanum in sputum samples using shotgun metagenomics on a benchtop sequencer. PeerJ. 2014;2014(1).

22. Goig GA, Cancino-Muñoz I, Torres-Puente M, Villamayor LM, Navarro D, Borrás R, et al. Whole-genome sequencing of Mycobacterium tuberculosis directly from clinical samples for high-resolution genomic epidemiology and drug resistance surveillance: an observational study. The Lancet Microbe. 2020;1(4).

23. Vasanthaiah S, Verma R, Kumar A, Bandari AK, George J, Rastogi M, et al. Culture-Free Whole Genome Sequencing of Mycobacterium tuberculosis Using Ligand-Mediated Bead Enrichment Method. Open Forum Infectious Diseases. 2024 Jul 1;11(7).

24. Brown AC, Bryant JM, Einer-Jensen K, Holdstock J, Houniet DT, Chan JZM, et al. Rapid whole-genome sequencing of mycobacterium tuberculosis isolates directly from clinical samples. Journal of Clinical Microbiology. 2015;53(7).

25. Doyle RM, Burgess C, Williams R, Gorton R, Booth H, Brown J, et al. Direct whole-genome sequencing of sputum accurately identifies drug-resistant mycobacterium tuberculosis faster than MGIT culture sequencing. J Clin Microbiol. 2018;56(8).

26. Nilgiriwala K, Rabodoarivelo MS, Hall MB, Patel G, Mandal A, Mishra S, et al. Genomic Sequencing from Sputum for Tuberculosis Disease Diagnosis, Lineage Determination, and Drug Susceptibility Prediction. Journal of clinical microbiology. 2023;61(3).

27. Jbara S, Herranz M, Sola-Campoy PJ, Rodríguez-Grande C, Chiner-Oms Á, Comas I, et al. Overlapping prison/community tuberculosis outbreaks in Costa Rica revealed by alternative analysis of suboptimal material. Transboundary and emerging diseases. 2022 May 1;69(3):1065–72.

28. Domínguez J, Acosta F, Pérez-Lago L, Sambrano D, Batista V, De La Guardia C, et al. Simplified model to survey tuberculosis transmission in countries without systematic molecular epidemiology programs. Emerging Infectious Diseases. 2019 Mar 1;25(3):507–14.

29. Genestet C, Perdigão J, Herranz M, Maus SR, Berland JL, Chiner-Oms Á, et al. Expanded tracking of a Beijing Mycobacterium tuberculosis strain involved in an outbreak in France. Travel medicine and infectious disease. 2021 Nov 1;44.

30. Acosta F, Agapito J, Cabibbe AM, Cáceres T, Sola C, Pérez-Lago L, et al. Exportation of MDR TB to Europe from Setting with Actively Transmitted Persistent Strains in Peru. Emerging infectious diseases. 2019 Mar 1;25(3):596–8.

31. Martínez-Lirola M, Jajou R, Mathys V, Martin A, Cabibbe AM, Valera A, et al. Integrative transnational analysis to dissect tuberculosis transmission events along the migratory route from Africa to Europe. Journal of travel medicine. 2021 May 1;28(4).

32. de Neeling AJ, Tagliani E, Ködmön C, van der Werf MJ, van Soolingen D, Cirillo DM, et al. Characteristic SNPs defining the major multidrug-resistant Mycobacterium tuberculosis clusters identified by EuSeqMyTB to support routine surveillance, EU/EEA, 2017 to 2019. Euro surveillance : bulletin Europeen sur les maladies transmissibles = European communicable disease bulletin. 2024 Mar 21;29(12).

33. Pérez-Lago L, Comas I, Navarro Y, González-Candelas F, Herranz M, Bouza E, et al. Whole Genome Sequencing Analysis of Intrapatient Microevolution in Mycobacterium tuberculosis: Potential Impact on the Inference of Tuberculosis Transmission. Journal of Infectious Diseases. 2014;209(1).

